# ARCH3D: A foundation model for global genome architecture

**DOI:** 10.64898/2026.02.23.707580

**Authors:** Nicholas Galioto, Cooper Stansbury, Alex Arkady Gorodetsky, Indika Rajapakse

## Abstract

Biological foundation models are transforming scientific discovery by creating information-rich representations that enable inference in low-data settings. Progress on these models has mainly been achieved by increasing input contextual information, e.g., base pairs or genes. Most work, however, focuses on DNA, RNA, and protein, leaving genome architecture, a fundamental component regulating processes like the cell cycle and cell-fate determination, underexplored. Here, we introduce ARCH3D: a foundation model for global genome architecture. ARCH3D uses a novel masked locus modeling task that increases input contextual information to include genome-wide contact profiles of loci spread across the entirety of the genome. We demonstrate this strategy captures global genome structure by showing ARCH3D embeddings preserve genomic spatial structure, reconstruct interchromosomal interactions under extreme sparsity, and enable identification of multi-way interactions. Ultimately, ARCH3D provides a potential structural foundation for building the virtual genome, an artificial intelligence-based model capable of simulating genome behavior and dynamics.

## Main

On the bleeding edge of the effort to facilitate mature biological machine learning (ML) has been the rapid development of various biological foundation models. A foundation model^1^ is meant to transform a dataset into a numerical representation (also known as an embedding) that can capture independent factors of variation within the data such that it can be used for a wide range of ML tasks with minimal adjustments or modifications. As examples, foundation models have been built to represent DNA sequence^2–6^, single-cell RNA-seq (scRNA-seq)^7–12^, and protein sequence^13–16^ data.

Much of the rapid progress in biological foundation models can be attributed to an incorporation of more contextual information into the embedding space, enabled by the transformer architecture^17^. In fact, recent research has shown that the performance of transformer-based biological foundation models is significantly elevated by extending the model’s context window. For example, the DNA sequence foundation models Evo2^13^ and AlphaGenome^3^ both outperformed their creators’ previous versions (Evo^4^ and Enformer^5^, respectively) by developing novel architectures that increase input sequence length to one million base pairs without loss of resolution. Similarly in transcriptomics, the scRNA-seq foundation model Geneformer^7,8^ showed improvement, in part, by increasing the context window from 2,048 to 4,096 genes.

While researchers continue to procure impressive leaps forward in the development of foundation models pertaining to the central dogma—DNA, RNA, and protein—less attention has been given to modalities measuring genome architecture. Chromatin organization plays a key role in important biological process such as gene transcription regulation^18^, DNA replication timing^19^, and cell-fate determination^20^. In one direction, researchers have sought to include structural information in foundation models by coupling chromatin accessibility data such as ATAC-seq^21^ or DNase-I^22^ with DNA sequence data. However, these one-dimensional assays do not capture spatial relationships between loci and are therefore insufficient for identifying spatially-localized phenomena between distal (including interchromosomal^23,24^) loci such as transcription factories^25^ and enhancer-promoter interactions^26,27^.

Currently, the most abundantly-available assay that measures genome-wide locus interactions is high-throughput chromosome conformation (Hi-C) data^28^. There has been a breadth of work applying machine learning techniques to enhance Hi-C analysis, most necessitating their own neural network-based encodings. Broadly speaking, these encoders can be divided into two categories: patch-based and locus-based approaches. Thus far, the context windows used for these encoders have been extremely restricted.

The patch-based approach uses a submatrix from the Hi-C map (usually from the diagonal) as input. This is the most common approach^29–32^, and indeed this is the approach used by the existing Hi-C foundation model HiCFoundation^33^. HiCFoundation is built upon a vision transformer architecture^34^ that takes a 224 × 224 submatrix and produces an embedding vector for every 16 × 16 patch within that submatrix. At the 5 kb pre-training resolution, this corresponds to a context window covering just 1.12 Mb of the genome. Moreover, this submatrix only measures the contact frequency of the 1.12 Mb region with itself, completely neglecting its interaction with the rest of the genome. The context window can of course be increased by coarsening the resolution, but this necessarily results in a loss of information. While the method has shown impressive utility for local analysis of Hi-C, it cannot account for interactions that occur at a genomic distance beyond the size of its input submatrix.

An alternative approach, used in models such as Hyper-SAGNN^35^, MATCHA^36^ and Higashi^37^, uses a selection of rows from the Hi-C matrix as input. Each row corresponds to a genomic locus, and the data in the row gives the observed contacts between that locus and all other loci in the genome. This input includes interactions that would otherwise be neglected by the patch-based approach, potentially offering a more global context window. However, the application of this approach has thus far been severely restricted in three main ways. First, in the three models listed previously, only data from intrachromosomal blocks are used as input. Second, input data is primarily limited to 1 Mb resolution. Third, the longest context window has included just five loci^36^, covering a small fraction of the genome. These restrictions reduce the input dimensionality and computational complexity but also prevent the model from learning higher-resolution and longer-range relationships within the Hi-C data.

In this work, we present ARCH3D, a foundation model for Hi-C capable of generating global representations of variable-length genomic loci using high-resolution Hi-C data from across the entirety of the genome. The model is built around a transformer backbone and trained with a novel variation of the celebrated masked language modeling task^7,38^ to produce information-rich embeddings encoding the global conformation of a given locus. The model uses a locus-based approach where each locus is input as the full Hi-C row, including interchromosomal regions, at 5 kb resolution. Additionally, we form the context window with loci selected from across the genome, thereby allowing the model to uncover relationships between distant loci. To ensure the model can still discern where each locus lies on the genome, we design a biologically-inspired positional encoding for the input. The random sampling approach has the added benefit of increasing the number of distinct input sequences combinatorially, significantly augmenting the training dataset to a scale suitable for a foundation model.

ARCH3D scales the locus-based encoding approach to a genome-wide level, making it well-suited for investigating long-range interactions and higher-order structures. We first confirm that the proposed masked locus modeling approach is effective in capturing global relationships by showing that the spatial structure of the low-dimensional embedding space preserves that of the genome in interphase nucleus. Then, we demonstrate that ARCH3D effectively leverages available data to accurately infer interchromosomal contacts, even when the data is downsampled to 99.58% sparsity in interchromosomal blocks. Lastly, we show ARCH3D embeddings enable identification of multi-way contacts from Hi-C data, achieving an average area under receiving operator curve of 0.930 across three cell lines, drastically outperforming the 0.787 value achieved by the state of the art. We give a summarized comparison of ARCH3D to the existing HiCFoundation^33^ in Table 1.

**Table 1.**
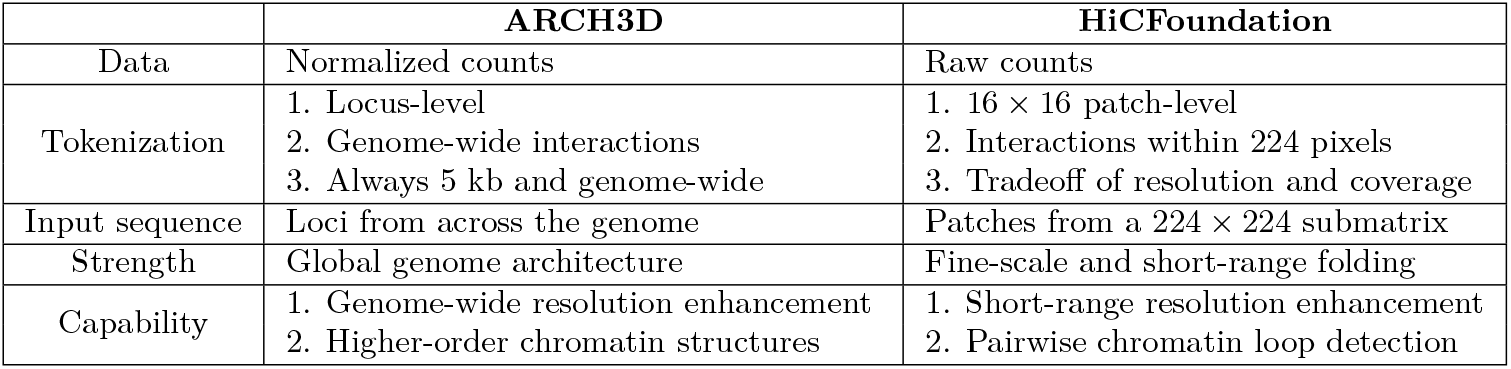
Comparison of ARCH3D and HiCFoundation^33^.

## Results

### Architecture and pre-training

ARCH3D is a chromosome conformation foundation model designed to produce locus-level embeddings of variable length that encode chromatin folding information, not just locally, but across the entirety of the genome. By producing embeddings that represent genomic loci rather than patches, ARCH3D outputs can be aligned directly with embeddings from DNA sequence and scRNA-seq foundation models, straightforwardly enabling multi-modal modeling of cellular processes. In this section, we describe our training corpus, model architecture design, and pre-training strategy.

For pre-training, we first assemble a training corpus consisting of all in-situ, dilution, and DNase bulk Hi-C data from the 4DNucleome and ENCODE consortia. Experiments with fewer than 10 million counts and those aligned with reference genomes other than GRCh38^39^ were excluded. In total, the pre-training set is comprised of 481 Hi-C experiments from 31 human tissues (Fig. 1f). The data were normalized with KR balancing^40^ and observed/expected normalization^28^, both at a 5 kb resolution. Then, pixel values were clipped at 15 to mitigate the effects of outliers on training. Lastly, 80% of the data was assigned to the training set and the remaining 20% was placed in the validation set.

**Fig. 1.**
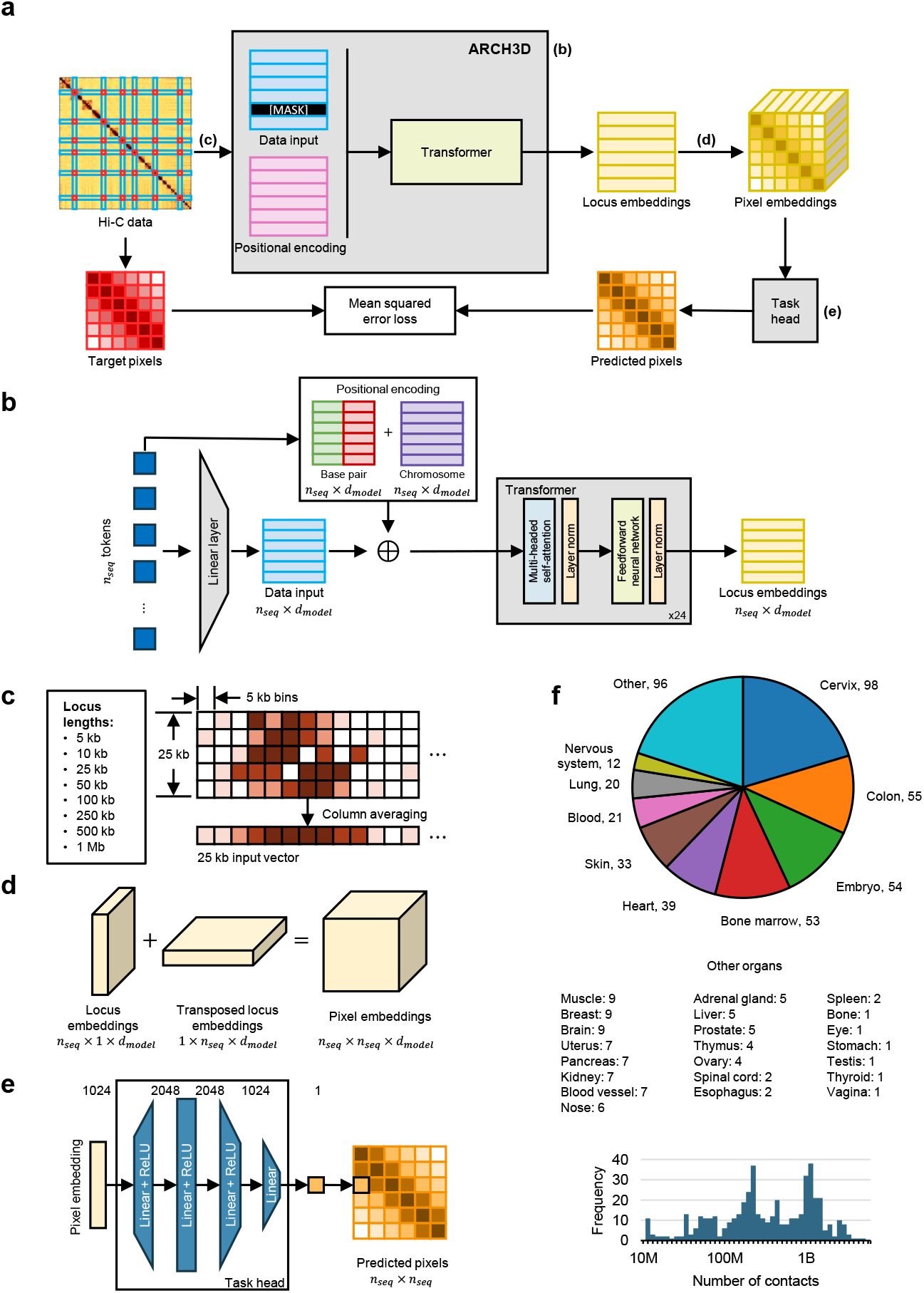
ARCH3D pre-training and architecture. **(a)** Overview of the masked locus modeling pre-training task. The Hi-C matrix is tokenized using *n*_*seq*_ = 1, 024 randomly-selected loci across the genome (represented as blue rectangles). To pre-train the model, 200 loci are masked, and a task head is used to predict pixel values between every pairwise combination of the selected loci (represented as red squares). The pre-training loss is the mean squared error between predicted and target pixels. **(b)** Architecture of ARCH3D. Tokens are projected into a low-dimensional space (*d*_*model*_ = 1, 024) using a linear layer. Then positional encodings comprised of the sum of base pair and chromosome encodings are added to the low-dimensional tokens. The result is passed through a transformer encoder with 24 layers. **(c)** Tokenization procedure. During pre-training, Hi-C data is provided at a 5 kb resolution, and the length of each locus is randomly selected from a pre-specified list of eight possible lengths. A locus longer than 5 kb is combined into a single vector by averaging along the columns of all rows lying within the locus. **(d)** Pixel embeddings. Every Hi-C pixel corresponds to a pair of genomic loci. Each pixel embedding is computed by summing the locus embeddings corresponding to that pixel’s locus pair. **(e)** Pre-training task head. The task head is a multi-layer perceptron consisting of three hidden layers and a linear output layer. The task head takes in a pixel embedding and returns the predicted value of that pixel. **(f)** Pre-training corpus. The pre-training data are sourced from 31 different human organs from the 4DNucleome and ENCODE consortia. Only data with at least 10 million contacts are used.

To facilitate the use of ARCH3D for biological modeling, we developed a novel tokenizer that yields locus-level tokens at variable lengths without sacrificing data resolution. For a given locus, the tokenizer returns the row from the Hi-C matrix corresponding to that locus. If the locus spans multiple Hi-C bins, then all the rows corresponding to bins intersecting with the locus are extracted, and the rows are averaged together to yield a single row vector (Fig. 1c). With this averaging step, the tokenizer is able to produce raw tokens corresponding to any genomic locus whose starting and ending boundaries are aligned with the boundaries of the 5 kb bins—the locus lengths are, of course, still limited by the binning of the Hi-C matrix at the base resolution. Allowing loci of varying lengths has the benefits of enabling direct alignment with other foundation models that tokenize along the genome and of increasing the total number of tokens in the training set to over two billion.

Once the sequence of raw tokens is generated, the tokens are embedded into the model’s latent space using a learnable linear layer. Given that the transformer encoder is a permutation-invariant transformation, positional information is not incorporated into the model’s transformation unless explicitly provided. We therefore develop a positional encoding composed of two parts to represent the position of a locus (Fig. 1b). The first part is a chromosome embedding in which each chromosome is represented by a learnable embedding vector. The other part is a base pair embedding derived from the sinusoidal encoding proposed by Vaswani et al.^17^. In this component, the beginning and ending base pairs are first divided by 10^6^ to map them into a range comparable with the encoding’s original formulation^17^. Then, the base pairs are passed through the sinusoidal encoding function to map each into one half of the model dimension. Lastly, the starting and ending encodings are concatenated together and added to the chromosome embedding to form the final positional encoding that gets added to the token embedding. The utility of each component is shown in Supplementary Fig. 1. The complete positional encoding is piecewise-continuous with discontinuities at the chromosomal boundaries, mimicking the physical piecewise-continuity of the genome.

Of note, this positional encoding encodes the global genomic positioning, rather than simply the positioning within a sequence. This property allows the input sequence to be formed from tokens located arbitrarily across the genome, not just sequentially, delivering two important benefits. First, using tokens from across the genome significantly increases the information contained in the sequence. As Hi-C measures whether two loci are in spatial proximity, adjacent loci naturally have very similar contact profiles, introducing redundancy into the input. By selecting loci that are farther apart, this redundancy is significantly mitigated. Furthermore, it allows the model to identify interactions at a range that is not constrained by the sequence length. The second benefit of using randomly-chosen loci is that it increases the set of distinct input sequences combinatorially, thereby serving as a data augmentation technique.

Lastly, we devised a masked locus modeling (MLM) task (Fig. 1a) that leverages self-supervised learning to ensure the model learns information-rich and context-aware representations of each locus. This task is an original variation of the masked language modeling task used to pre-train BERT^38^. In MLM, we build an input sequence composed of 1,024 randomly selected loci, where each locus length is randomly chosen from the set of 5 kb, 10 kb, 25 kb, 50 kb, 100 kb, 250 kb, 500 kb, and 1 Mb. From this sequence, 200 randomly-chosen loci are replaced with a learnable [MASK] vector, and the rest are passed through an initial linear layer. Then, positional encodings are added to both the masked and unmasked loci, and the sequence is passed through a transformer encoder with 24 layers. The architecture of the transformer encoder^17^ follows that of the BERT-large model^38^ and enables the model to discover relationships between distal loci in the input sequence. At the output, each unordered pair of loci is combined into a pairwise embedding (Fig. 1d) and passed through a multi-layer perceptron (MLP) task head that predicts the pixel value corresponding to that locus pair (Fig. 1e). These predictions are generated for all pairs regardless of whether or not they contain a masked locus. Including the unmasked loci in the pre-training objective ensures the model learns to represent the masked and unmasked loci similarly. This is important to mitigate the discrepancy between pre-training and inference, in which only unmasked loci are used.

### ARCH3D embeddings preserve the spatial arrangement of chromatin in interphase nucleus

To understand how ARCH3D organizes loci within its embedding space, we examined distances between embedding vectors. For this experiment, we considered cell lines H1-hESC, GM12878, and IMR-90 at 100 kb resolution. To generate the ARCH3D embeddings, we partitioned the genome into 100 kb loci and randomly placed them into sequences of length 1,024. Once fewer than 1,024 loci were left, the remaining loci were added round robin into the sequences until all loci had been assigned to a sequence. Then, each sequence was passed through ARCH3D to generate an embedding of every locus.

Next, we computed the Euclidean distance between every pair of embedding vectors. We then grouped the distance values according to the chromosome pair from which each originated, and Fig. 2a shows the average distance within each of these groups. We first notice that the intrachromosomal blocks have the lowest average distance within all cell lines, indicating that embeddings from the same chromosome tend to be positioned closer than embeddings from different chromosomes. This intrachromosomal clustering within embedding space mirrors the chromosome territories within the nucleus^41^.

**Fig. 2.**
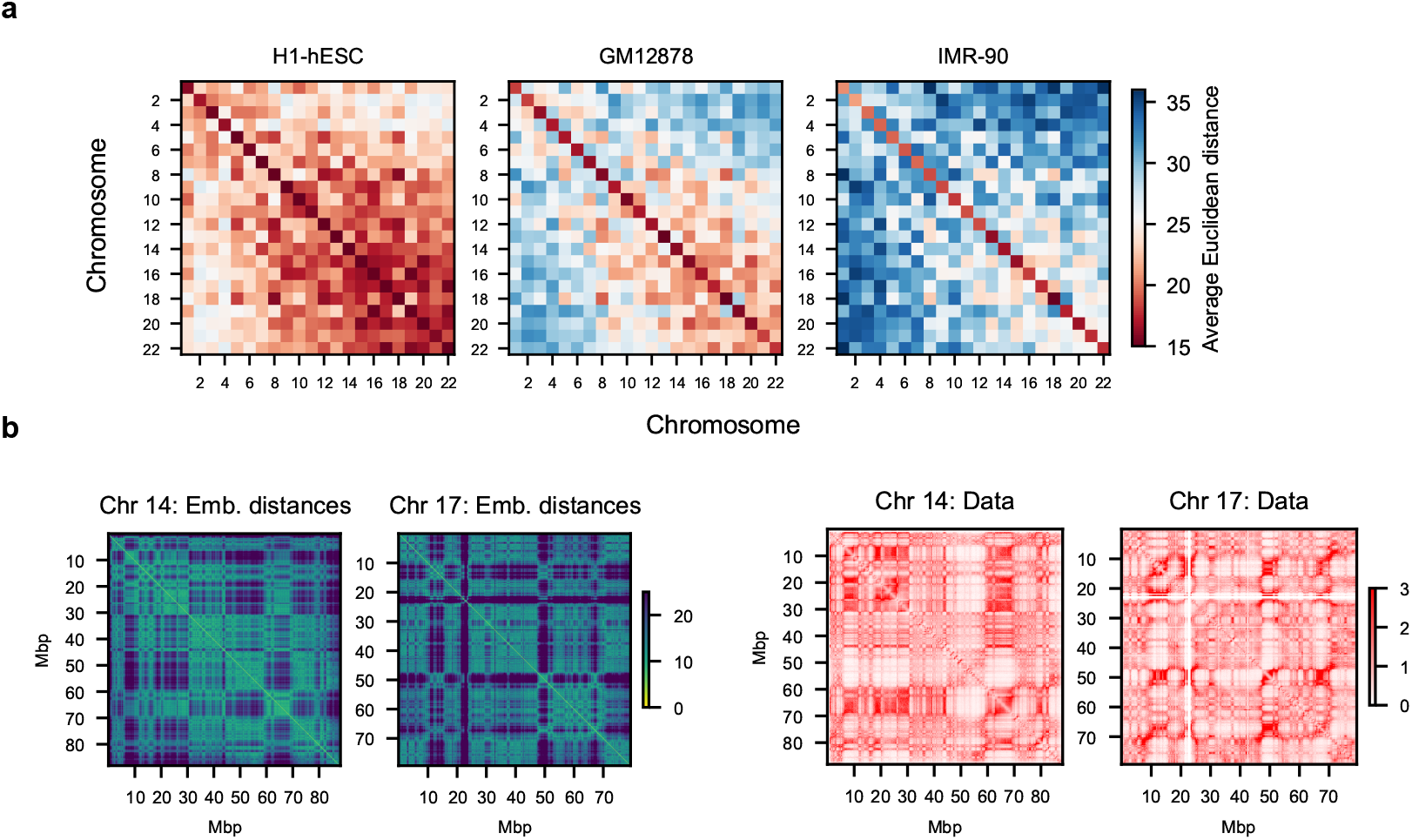
Spatial structure of ARCH3D embedding space. **(a)** Average inter- and intrachromosomal distances across three cell types. Each pixel belongs to a chromosome pair, and the pixel value is the average Euclidean distance between embeddings of loci lying on either of the paired chromosomes. We observe that (1) embeddings from the same chromosome are positioned more closely together than embeddings from different chromosomes, reflecting the existence of chromosome territories. (2) Embeddings from smaller chromosomes tend to be in closer proximity than embeddings from larger chromosomes, mirroring the interaction preferences of the physical genome^28^. (3) The more differentiated the cell type, the farther apart embeddings are spaced. Again, this is in agreement with findings on the spatial arrangement of chromosomes in the nucleus^42^. **(b)** Comparison of intrachromosomal embedding distance maps (left) and Hi-C contact maps (right) of chromosomes 14 and 17 for IMR-90. The compartmentalization patterns of the embedding space mirror those of the Hi-C map, indicating that structural information from the Hi-C data is preserved in the ARCH3D embedding space.

We further notice that embeddings in H1-hESC are overall positioned more closely together than in GM12878, whose embeddings are in turn positioned more closely than in IMR-90. Notably, this ordering matches the developmental lineage of the cell types: H1-hESC is less differentiated than GM12878, which is less differentiated than IMR-90. As a cell becomes increasingly differentiated, more of its chromatin closes and fewer interchromosomal contacts are observed^42^. Fig. 2a shows that this developmental phenomenon is reflected in the embedding distances. Moreover, the interchromosomal blocks that have lower average distances tend to come from the smaller, more gene-rich chromosomes. An analogous observation has been made for Hi-C contacts where smaller chromosomes show higher interchromosomal interactions than their larger counterparts^28^.

In addition to mirroring the interchromosomal relationships of the physical genome, we show in Fig. 2b that structure is also mirrored intrachromosomally. Fig. 2b compares the intrachromosomal embedding space of chromosomes 14 and 17 to the corresponding Hi-C maps at 100 kb resolution. We observe very similar compartmentalization patterns, indicating that much of the structural information of the Hi-C data has been preserved in the low-dimensional embedding space. Overall, the organization of the embedding space mirrors that of the nucleus, illustrating how ARCH3D embeddings capture the global structure of the genome.

### ARCH3D infers distal contacts under extreme sparsity

The sparsity and noisiness of bulk Hi-C can vary substantially depending on sequencing depth/coverage and experimental procedures. At lower sequencing depths and coverages, the sparsity and noise increases, and the quality of the Hi-C data suffers. This degradation is especially prominent as the distance between loci increases and contact frequencies rapidly decrease. The quality of interchromosomal regions suffers the worst of these effects. Because interchromosomal interactions typically occur at greater distances than intrachromosomal ones^43^, they are normally more difficult to detect with ligation-based assays such as Hi-C, and low sequencing depth only exacerbates this issue.

Promising research has shown that many of the high-coverage contact patterns along the Hi-C diagonal can be reliably reconstructed from low-coverage data, a task often referred to as resolution enhancement. Models such as HiCFoundation^33^, HiCARN^30^, HiCSR^31^, and HiCNN^32^ have all demonstrated impressive abilities to upscale Hi-C to a coverage level at least 16x greater than the input data. However, all of these models use a patch-based approach where missing features are inferred using available data contained within a relatively small neighborhood of the imputed region. While existing models have proven effective under this setting, the approach cannot be extended to regions where little to no data are available.

ARCH3D’s global approach to locus-level representation promises to dramatically extend the domain of resolution enhancement to regions thus far unreachable by existing methods. The masked locus modeling scheme trains ARCH3D to leverage all available contact information from a given locus to infer its interaction frequencies across the entire genome. As a pre-trained model, however, ARCH3D’s prediction quality is limited by the noise and sparsity encountered in the pre-training corpus. To accurately predict the observed pixel values during pretraining, ARCH3D needed to account for the individual noise/sparsity profiles of each experiment. To overcome this limitation, ARCH3D must be fine-tuned to infer high-coverage contact maps regardless of the input data sparsity.

In this procedure, we gathered eight Hi-C experiments from the highest-coverage dataset available^44^. We downsampled each experiment to 10% and 1% of its full number of reads. At 10% and 1% coverage, the highest-depth experiment (≈ 4B contacts) has fewer contacts than 46% and 91% of the pre-training corpus, respectively. We then ran a slightly-modified (Methods) version of the pre-training task using the 1%, 10%, and 100% data as inputs and only the corresponding 100% data as targets. Six of the experiments were included in the training set and two were withheld for testing.

After training, we tasked the model with inferring contact profiles of the two test experiments and compared the results with HiCFoundation’s resolution enhancement model^33^. Fig. 3a shows predicted intrachromosomal contacts at 100 kb resolution on the IMR-90 cell line (≈ 1B counts) from ARCH3D and HiCFoundation^33^. At 1% coverage, the input data has only 10M counts, placing it right at the lower limit of the pre-training corpus. Predictions on HUVEC (≈ 131M counts) are located in Supplementary Fig. 2. We observe two main differences in reconstruction from ARCH3D and HiCFoundation that highlight key distinctions between the foundation models.

**Fig. 3.**
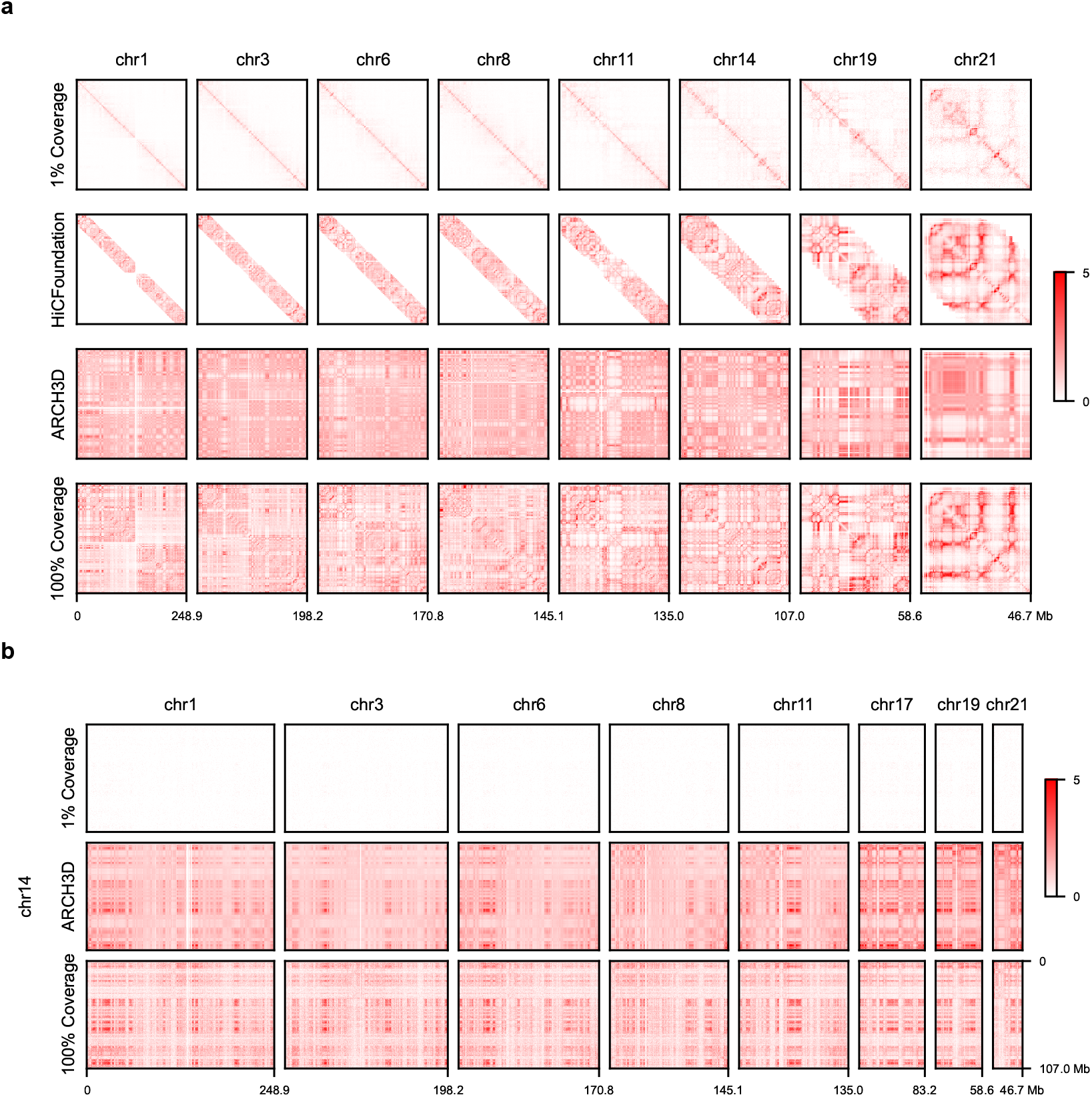
Resolution enhancement on IMR-90 at 100 kb resolution from 1% coverage (≈ 10M counts). **(a)** Prediction of intrachromosomal contacts. HiCFoundation^33^ shows strong prediction of fine-scale patterning along the diagonal, but it does not predict interactions between loci more than 22.4 Mb apart. In contrast, ARCH3D can predict across the entirety of the contact map, not just the diagonal. While these reconstructions are not as detailed as HiCFoundation’s they still capture much of the compartmentalization structure of the chromosomes. **(b)** Prediction of interchromosomal contacts. In the interchromosomal regions, the downsampled data are incredibly sparse (0.42% of pixels are nonzero), rendering HiCFoundation unusable. Conversely, ARCH3D leverages data from other regions to accurately infer the interchromosomal interactions. This is illustrative of the tradeoff between capturing distal interactions and fine-scale, near-diagonal patterns.

First, we observe that HiCFoundation excels at preserving the fine-scale features within the data compared to ARCH3D. This is explained by the differences in the target data used for training. In HiCFoundation, the model is tasked with reconstructing a 224 × 224 pixel submatrix. For resolution enhancement specifically, these submatrices are centered on the Hi-C diagonal, meaning that they are the most densely-sampled submatrices of the entire Hi-C map. The result is that the data used to fine-tune HiCFoundation have minimal noise and sparsity. In contrast, the data entering into the loss used to train ARCH3D come from pixels farther from the diagonal, including many from interchromosomal blocks, that have only a small number of observed counts and high sparsity. Consequently, the terms in the loss function that capture the error in reconstructing fine-scale, near-diagonal features are dominated by the noise from distal contacts. This prevents ARCH3D from learning to resolve the more detailed patterning near the diagonal.

The second observation is that ARCH3D is capable of making predictions anywhere within the chromosomal contact maps while HiCFoundation is restricted to a window of 224 pixels along the diagonal. This observation illustrates the tradeoff between capturing fine-scale details along the diagonal and being able to infer long-range interactions. To further elucidate this point, we plot several interchromosomal blocks in Fig. 3b. At 1% coverage, the interchromosomal regions are so sparse that only 0.42% of pixels are nonzero. Regardless, ARCH3D leverages the available contact data from other regions to accurately infer the interchromosomal interactions. This gives ARCH3D a marked advantage over patch-based models, which have no method of transferring information across distant regions in the Hi-C map.

### ARCH3D identifies higher-order structures from pairwise data

The next generation in chromosome conformation capture is Pore-C^48^, an extension of Hi-C that uses long-read sequencing to measure spatial interactions of two or more chromatin segments. Observing higher-order interactions gives greater insight into genome architecture because it reveals whether certain interactions occur cooperatively or competitively. Indeed, Pore-C has led to the detection of multi-locus hubs associated with increased transcriptional activity^49^ and histone locus bodies^50^. However, Pore-C data is extremely limited in comparison to the availability of Hi-C data due to its greater cost and dependence on long-read sequencing. We therefore seek to leverage deep learning and ARCH3D to predict higher-order chromatin interactions from Hi-C.

To begin, we created a corpus of Pore-C data at 100 kb resolution using GM12878 data from Deshpande et al.^48^ and BJ and IR fibroblast data from Dotson et al.^49^ For practicality reasons, we considered only 3-way, 4-way, and 5-way contacts. In total, this yielded nearly 90 million contacts, 72% of which span multiple chromosomes (Supplementary Table 4). Clearly, effectively leveraging interchromosomal interactions is key for this task, and patch-based methods such as HiCFoundation^33^ are not applicable here.

We next expanded the multi-way Pore-C data into pairwise data known as “virtual Hi-C”^50^ (Methods). We composed the training set of 100,000 unique hyperedges from each cell type and reserved the remaining data for testing. The training data represent 0.24%, 15.59%, and 38.54% of the total number of unique hyperedges for GM12878, BJ fibroblast, and IR fibroblast, respectively. To train, we froze the ARCH3D encoder and trained a task head composed of a four-layer transformer encoder and a linear output layer to predict whether or not a set of loci comprised a hyperedge. The training scheme is depicted in Fig. 4a and detailed in Methods.

**Fig. 4.**
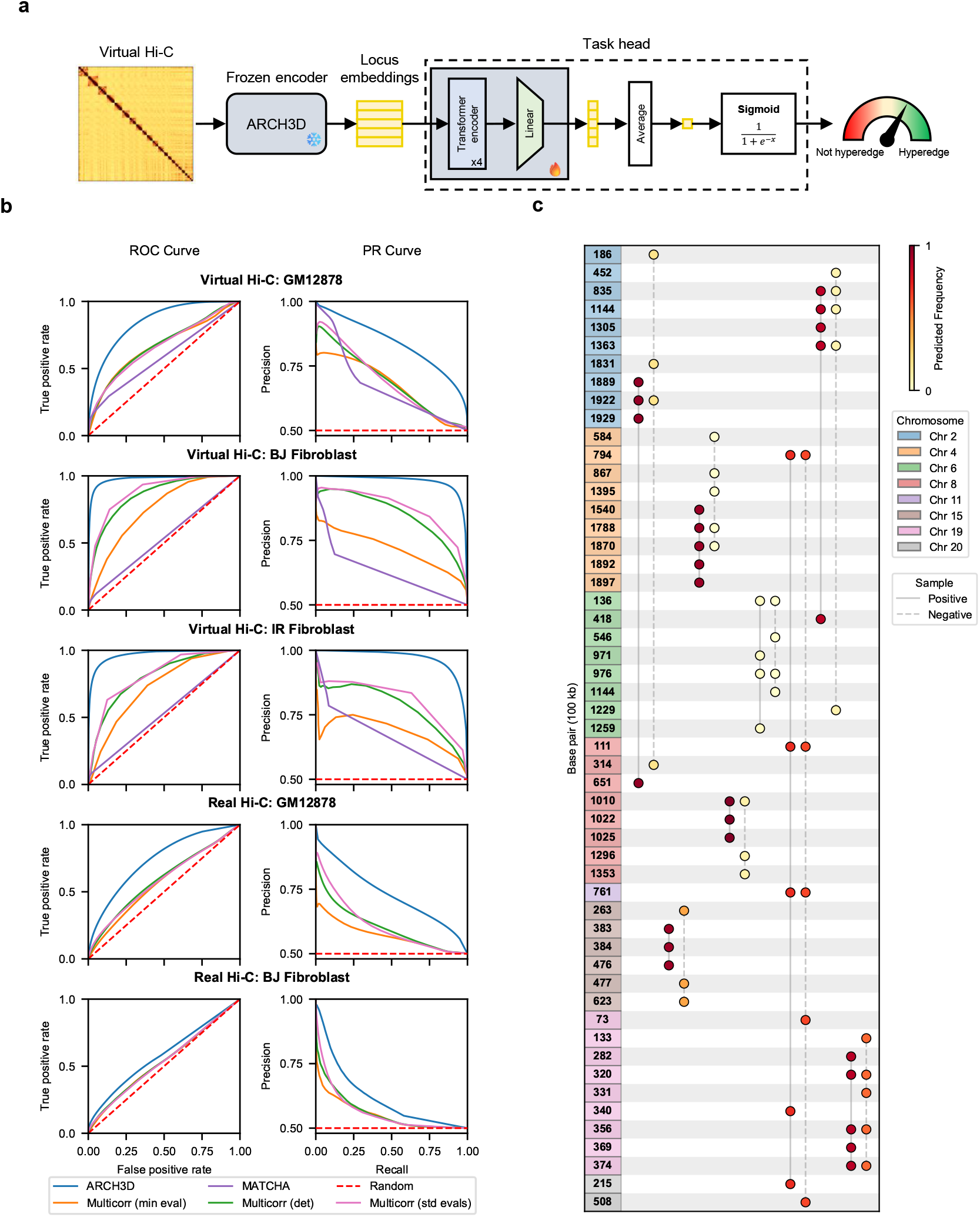
Hyperedge prediction. **(a)** The training scheme. Pore-C data is expanded into virtual Hi-C that is then fed into the pre-trained ARCH3D to generate locus embeddings. The embeddings corresponding to loci in the candidate hyperedge are passed into a trainable task head that predicts the probability that the loci comprise a hyperedge. **(b)** Comparison of ROC and PR curves. The left (right) column gives the ROC (PR) curve, and each row shows a different input dataset. ARCH3D shows significant improvement over all baselines. **(c)** Randomly-selected positive and negative hyperedges from GM12878. The circles in each column indicate the nodes in a hyperedge, and the style of the line (solid or dashed) denotes whether or not the hyperedge was observed in the Pore-C dataset. The negative samples are generated from the positive samples positioned directly to their left using the procedure from MATCHA^36^. The predicted probability that a hyperedge is observed is denoted by the color of the circles in the hyperedge.

As comparison methods, we considered the deep learning model MATCHA^36^ and three statistical methods for computing multi-correlations. Because MATCHA makes predictions using only node indices, it cannot be trained across multiple cell types at once. We accommodated this limitation by training MATCHA from scratch on each cell type separately. For each cell line, MATCHA used 20% of the total unique hyperedges to train. The multi-correlation coefficients were computed using the correlation matrix between all nodes in each candidate hyperedge. The three methods are defined in terms of the minimum eigenvalue^45^, determinant^46^, or standard deviation of eigenvalues^47^ of this correlation matrix (Eqs. (16)–(18) in Methods).

We tested the approaches in two settings. In the first setting, we used virtual Hi-C as input. This is the setting considered by MATCHA, and it serves to test how well each model can predict higher-order interactions from ground truth pairwise interactions. In other words, this setting does not consider any mismatch between the pairwise and multi-way interaction data. In the other setting, we used real Hi-C data as input. This setting is more consistent with a real-world setting in which Pore-C data are not always available. As IR fibroblast Hi-C is unavailable, we excluded the cell line from this setting.

To evaluate each method, we computed the receiving operator curves (ROC) and precision-recall curves (PR). The curves are shown in Fig. 4b and the area under each curve is given in Table 2. In the figure, we plotted the curves given by random guessing as red dashed lines. ARCH3D outperforms all baselines across all cell lines and testing settings. Compared to GM12878, ARCH3D shows slightly improved performance on the BJ and IR fibroblasts, but this is likely because a greater percentage of the fibroblast datasets were included in the training set. We also observe that ARCH3D gives the best performance on the real Hi-C data, indicating that the model generalizes well to unseen datasets.

**Table 2.** Summary of hyperedge prediction results. ARCH3D was trained on 100,000 samples from each cell line, representing 0.24%, 15.59%, and 38.54% of the total GM12878, BJ fibroblast, and IR fibroblast datasets, respectively. Testing is conducted on the virtual Hi-C created from the Pore-C dataset and real Hi-C taken directly from separate Hi-C experiments. MATCHA cannot be applied to a new dataset without being re-trained from scratch, so we cannot test it on real Hi-C. Across all datasets and testing settings, ARCH3D shows the best performance.

| Method | Metric | GM12878 |  | BJ Fibroblast |  | IR Fibroblast |
| --- | --- | --- | --- | --- | --- | --- |
|  |  | Virtual | Real | Virtual | Real | Virtual |
| ARCH3D | AUROC | **0.848** | **0.737** | **0.976** | **0.584** | **0.966** |
|  | AUPR | **0.839** | **0.731** | **0.978** | **0.618** | **0.968** |
| MATCHA | AUROC | 0.587 | N/A | 0.536 | N/A | 0.531 |
|  | AUPR | 0.668 | N/A | 0.627 | N/A | 0.664 |
| Multi-correlation<br>(Min. eigenvalue<sup>45</sup> Eq. (16)) | AUROC | 0.665 | 0.573 | 0.751 | 0.538 | 0.725 |
|  | AUPR | 0.673 | 0.571 | 0.717 | 0.552 | 0.685 |
| Multi-correlation<br>(Determinant<sup>46</sup> Eq. (17)) | AUROC | 0.672 | 0.592 | 0.859 | 0.540 | 0.801 |
|  | AUPR | 0.687 | 0.601 | 0.850 | 0.560 | 0.783 |
| Multi-correlation<br>(Eigenvalue std.<sup>47</sup> Eq. (18)) | AUROC | 0.664 | 0.586 | 0.883 | 0.538 | 0.815 |
|  | AUPR | 0.688 | 0.607 | 0.874 | 0.562 | 0.809 |

**Table 3.** Pre-training hardware summary. The runs comprising stage 2 used either 16 GPUs and 3.2 TB or the full four nodes comprising 32 GPUs and 4 TB.

| Stage | Nodes | GPUs | RAM (TB) | Epochs | Time (h) | GPU hours |
| --- | --- | --- | --- | --- | --- | --- |
| 1 | 1 | 8 | 1 | 6,000 | 504 | 4,032 |
| 2 | 4 | 16/32 | 3.2/4 | 5,700 | 384 | 9,600 |

Lastly, we plotted a visualization of a small number of hyperedge predictions using an incidence matrix in Fig. 4c. This style of visualization of hyperedges comes from work on hypergraphs^51,52^. On the y-axis are 100 kb bins, colored by the chromosome to which they belong, and each column represents a unique hyperedge. As an example, the first column shows a hyperedge consisting of nodes at 1.889, 1.922, and 1.929 Mbp on chromosome 2, and a node at 0.651 Mbp on chromosome 8. This hyperedge was observed in the Pore-C dataset, so the nodes are connected by a solid line. The column directly to the right shows a hyperedge with nodes at 0.186, 1.831, and 1.922 Mbp on chromosome 2 and another node at 0.314 Mbp on chromosome 8. This hyperedge does not appear in the Pore-C, so it is connected by a dashed line. The model predictions are indicated by the color of the nodes within each hyperedge where dark red (light yellow) denotes a high (low) frequency of observance of that hyperedge. ARCH3D correctly predicts that the first hyperedge will be observed much more frequently than the second. Although other predictions are more ambiguous, ARCH3D still drastically outperform existing baselines on this dataset, as evidenced by the ROC and PR curves.

## Discussion

In this work, we presented ARCH3D, a chromosome conformation foundation model for global genome architecture. ARCH3D creates representations of genome structure through locus-level tokenization and a novel masked locus modeling pre-training scheme. We showed that ARCH3D structures its embedding space in a way that naturally mirrors that of the physical genome, reflecting a preservation of the Hi-C data’s structural information. We then demonstrated that ARCH3D’s global approach to representation learning enables it to significantly extend the task of resolution enhancement to regions of extreme sparsity that were previously unreachable by existing, patch-based methods. This included faithfully reconstructing interchromosomal contacts from Hi-C experiments with as little as 0.42% nonzero pixels. Finally, we showed that ARCH3D embeddings could be used to reconstruct Pore-C data from Hi-C at 100 kb resolution, achieving an average AUROC value of 0.930, a significant improvement over both the 0.551 value given by MATCHA^36^ and 0.787 value given by the best multi-correlation method^47^.

There are several exciting directions for future work. In one direction, ARCH3D can be combined with other data modalities to enrich the representation quality of both itself and other foundation models. For example, incorporating ATAC-seq, DNaseI, and/or methylation profiles from ChIP-seq data with ARCH3D would lead to more complete representations of epigenetic information within a cell. Moreover, ARCH3D could be trained to produced embeddings conditioned on single-cell embeddings from other models, allowing it to better capture cell heterogeneity. This extension to single-cell data would lead to stronger integration with single-cell foundation models of other modalities. For example, single-cell ARCH3D embeddings could serve as positional encodings for transformer-based transcriptomic models^7^.

In another direction, ARCH3D can be used in conjunction with other foundation models to model complex phenomena. Modeling biological processes with foundation models is still an emerging area. Presently, the single notable example is State^53^, which uses embeddings from the protein language model ESM-2 within a greater neural network-based model to predict cellular perturbation response. Similarly, ARCH3D, or any of its extensions discussed in the previous paragraph, could act as building blocks when modeling processes that depend heavily upon genome folding and epigenetics. Specifically, we envision a virtual genome that can enable in silico perturbation experiments to help identify optimal strategies for cell reprogramming. Such a model would integrate the genome’s structure using ARCH3D embeddings and its genetic activity using transcriptomic foundation models. Once constructed, the virtual genome could efficiently identify highly-valuable experiments to be conducted in the wet lab, accelerating the pace of science. Further, pairing the virtual genome with recent advances in self-driving labs could even lead to fully-automated biological discovery.

Overall, ARCH3D introduces a representation of global 3D genome architecture to an ecosystem of biological foundation models that have so far largely focused on DNA, RNA, and protein. While ARCH3D and other foundation models display impressive performance individually, intelligent fusion of these modalities will lead to remarkable capabilities.

## Methods

### Data

The pre-training data are in-situ, dilution, and DNase Hi-C downloaded from the 4DNucleome^1^ and ENCODE^2^ consortia. We filtered out all experiments that either were not aligned to the GRCh38 reference genome or did not have at least 10 million reads..

We normalized every Hi-C matrix with KR balancing^40^ and observed/expected normalization^28^. The KR balancing serves to remove technical artifacts and map experiments of different read counts into the unit interval, and observed/expected normalization removes the skew toward diagonal entries in the data. If any pixels are greater than one after KR balancing, we set them to be zero. To handle interchromosomal contacts during observed/expected normalization, we divide pixels in the interchromosomal regions by the average of all non-zero interchromosomal pixels within that experiment. Both of these normalization steps were performed at 5 kb resolution. Lastly, to mitigate the effects of outliers, we clip all pixels to the interval [0, 15].

### ARCH3D architecture

As the backbone of ARCH3D, we use the BERT-large encoder architecture^38^. It is composed of 24 transformer layers, each with hidden dimension *d* = 1, 024, feedforward dimension of 4,096, 16 attention heads, and ReLU activation functions. This encoder contains 302 million learnable parameters. Since transformer models require the input be formulated as a sequence, we must devise a method for transforming Hi-C data into an amenable format. To this end, we require a tokenization scheme, a method of representing each token numerically, and a positional encoding.

#### Notation

To begin, we introduce some notation. Let ℝ be the set of real numbers, ℤ_≥0_ be the set of non-negative integers, and ℕ be the set of positive integers. We denote matrices as uppercase and bold **A**, vectors as lowercase and bold **a**, and scalars as lowercase and italicized *a*. A genomic locus that begins on the *a*th base pair of chromosome *c* and ends on the *b*th base pair (exclusive) of chromosome *c* is denoted as 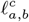. Here, *a < b* ∈ ℤ_≥0_ and *c* ∈ 𝒞:= {0, …, 21}. In this definition, each locus cannot belong to more than a single chromosome. The interval containing the entire chromosome *c* is denoted as *ℓ*^*c*^, and the length of the chromosome is denoted as | *ℓ*^*c*^ |. Note that all indexing here is zero-based.

#### Tokenization

When using a transformer architecture, the data must decomposed into a sequence of discrete elements known as tokens^54^. Since transformers are specifically designed to find relationships between tokens within a sequence, the choice of tokenization scheme has a direct impact on the type of relationships the model can identify. As we are interested in accounting for the relationship in conformation between loci, we elect to tokenize the Hi-C data at the locus level. Furthermore, we would like a tokenization scheme flexible enough to represent any locus of interest on the genome, such as genes, promoters, enhancers, etc., regardless of locus length.

To achieve tokenization of loci at arbitrary lengths, we first consider tokenizing loci at a single, short length and then extending to longer lengths. Consider the set of loci spanning *r*_∗_ base pairs, defined by the basis set *B*_*r*∗_ as

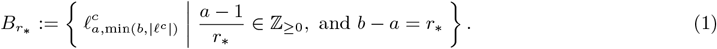

Here, we have constrained the loci to be non-overlapping intervals beginning from the start of each chromosome to be consistent with the binning scheme used in Hi-C data.

From this basis *B*_*r*∗_, we construct a domain *D* ⊃ *B*_*r*∗_ as the set of all loci in *B*_*r*∗_ and the unions of adjacent basis elements. This set can be written as

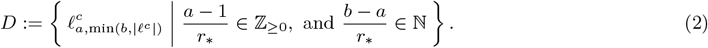

We consider each element of *D* to be a valid token. Again, this is consistent with how Hi-C data is binned at varying resolutions.

#### Numerical representation

With a tokenization scheme defined, we must now be able to represent each token numerically. Let 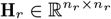 be the normalized Hi-C matrix where the number of bins *n*_*r*_ depends on resolution *r* by the relationship *n*_*r*_ = ∑_*c*∈𝒞_ ⌈|*ℓ*^*c*^|/*r*⌉, where ⌈· ⌉ is the ceiling operator. In this work, we use the base resolution *r*_∗_ = 5, 000 because it is the finest resolution shared across the pre-training dataset. With GRCh38^39^ as the reference genome, this yields *n*_*r*∗_ = 575, 010. The bin indices for a locus 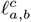 are given by the function

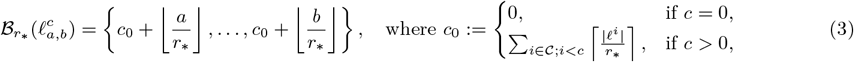

where ⌊·⌋is the floor operator.

We denote the numerical representation of a locus 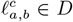 as the vector 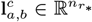. Each entry in 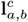 is a non-zero column average defined as

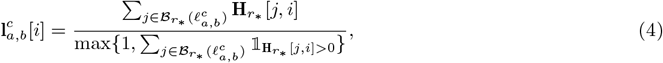

where 𝟙 is the indicator function. Given that the transformer requires each token to lie in the hidden space ℝ^*d*^, we create a linear input layer with weights 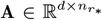 and bias **b** ∈ ℝ^*d*^ that projects 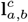 into ℝ^*d*^ via the transformation 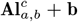. Strategies for reducing the number of parameters for this input layer are discussed in Supplementary Note 1.

#### Positional encoding

To complete the input embedding of a locus, we must define a function *γ*: *D* ↦ ℝ^*d*^ that encodes the locus’s genomic positioning. In transformer models for language and images, the position encoding only represents the token’s position within the sequence, but such an approach cannot capture the varying locus lengths represented by the tokens, nor the increased distance between loci on different chromosomes. To address these shortcomings, we develop separate encodings for the base pairs and chromosomes.

For the base pair encoding, we encode the first and last (inclusive) base pairs on the locus using the sinusoidal positional encoding *ρ*: ℝ _≥0_ ↦ [0, 1]^*d*/2^ from^17^ and concatenate them together. Given a base pair position *x*, the encoding is defined as

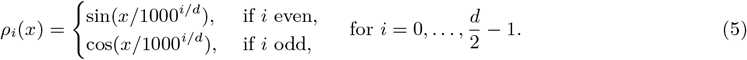

For the chromosome encoding, we create a codebook **C**: 𝒞 ↦ ℝ^*d*^ containing a learnable vector for each chromosome. The full position encoding is simply the sum of these two encodings

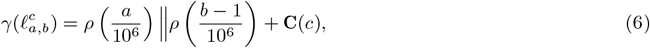

where ‖ denotes the concatenation operator. Note that we have divided the base pair coordinates by 10^6^ to map them into a comparable magnitude as the original transformer model^17^. We now have the final input token embedding of a locus 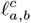 defined as

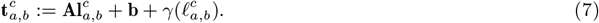

### Pre-training

In this section, we describe the methodology for pre-training ARCH3D. This description includes how the input sequence is formed, the architecture of the task head, the novel masked locus modeling task, and the optimization scheme.

#### Sequence formation

The transformer encoder uses data from across the input sequence to enrich the returned embeddings. The more context supplied in the input sequence, the more interactions can be identified between tokens to improve the learned representations. In the ideal case, the full Hi-C matrix would be supplied to the model, but due to the size of the matrix, this is computationally infeasible. One solution is to lengthen the input loci to cover more of the genome for a given sequence length, but this runs the risk of obscuring fine-scale features in the data. Instead, we make the observation that the contact profiles of adjacent loci are very similar and therefore introduce redundant information into the input sequence. In an effort to maximize the information contained in the sequence, we elect to form the input sequence with loci randomly selected from across the genome. Because the biologically-informed positional encodings from the previous section describe genomic, rather than sequence, positioning, we are not restricted to forming the sequence from contiguous loci like other biological foundation models. Moreover, we fill the sequence with loci of different lengths to increase coverage of the genome and ensure that the model performs well at any length. To balance the tradeoff between coverage and resolution, we randomly sample the locus lengths from a set that includes a range of lengths at varying levels of genome coverage and fine-scale detail.

To describe the sampling procedure, it is useful to think of a locus as a set of contiguous row indices, i.e., slices, of the Hi-C matrix. The goal is to randomly pick a set of *N* non-overlapping slices 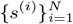 that will serve as the input sequence. Each slice *s*^(*i*)^ in this set consists of *λ*^(*i*)^ ∈ ℕ bins. Let 𝒰 {*S*} denote the uniform distribution over a bounded set *S*. We first sample the number of bins in each slice from a pre-specified set *λ*^(*i*)^ ∼ 𝒰 {1, 2, 5, 10, 20, 50, 100, 200} for *i* = 1, …, *N*. Given that nearly all of the bins are of resolution 5 kb (the only exceptions are bins at the ends of chromosomes), this roughly corresponds to lengths of 5 kb, 10 kb, 25 kb, 50 kb, 100 kb, 250 kb, 500 kb, 1 Mb.

Next, we position these slices on the Hi-C matrix without overlap by randomly selecting how many bins should separate each slice. First, we compute the remaining number of bins 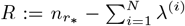 and partition them into *N* + 1 subsets. This is done by first sampling *u* ^(*i*)^∼ 𝒰 [0, 1] for *i* = 1, …, *N* + 1. We require that the samples be integers summing to *R*, so we transform them into values 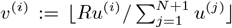 for *i* = 1, …, *N* and 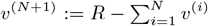. We then set the *i*th slice to begin at the bin 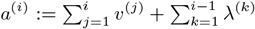 and ending at the bin *b*^(*i*)^ = *a*^(*i*)^ + *λ*^(*i*)^ (exclusive). This ensures that the slices do not overlap and do not go beyond the total number of bins. Lastly, any slice that crosses onto a second chromosome is truncated to lie only on a single chromosome while retaining the majority of its bins. The result is a sequence of slices *s*^(*i*)^ = {*a*^(*i*)^, …, *b*^(*i*)^ − 1} for *i* = 1, …, *N* corresponding to a set of genomic loci 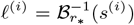, where 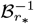 represents the inverse of the function ℬ_*r*∗_ defined in Eq. (3). Each locus *ℓ*^(*i*)^ has the numerical representation **l**^(*i*)^ as defined in Eq. (4).

Next, we describe the masking procedure. Let 𝒯= {1, …, *N*} be the index set of a given sequence. The set of masked tokens is a non-empty set ℳ ⊂ 𝒯 of the indices of tokens to be replaced by a learnable [MASK] vector **m**. When pre-training ARCH3D, the set ℳ was formed by randomly sampling 200 indices from 𝒯 without replacement. Then, the numerical form of each token in a sequence is defined as

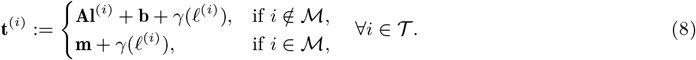

The sequence 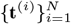 serves as the input to the transformer backbone.

#### Task head

At the encoder output, we first add every unordered pair of embeddings together to get *N* (*N* + 1)/2 pairwise embeddings. Each of these pairwise embeddings is transformed through a task head to predict the corresponding pixel value. As a task head, we use an MLP with three hidden layers and a linear output layer specified as

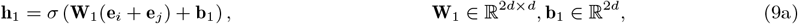

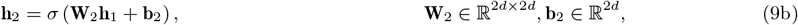

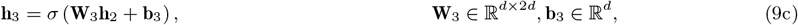

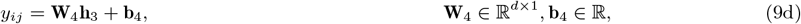

for any *i, j* ∈ 𝒯, such that *i* ≤ *j*. Here **e** ∈ ℝ^*d*^ is a locus embedding, **W**_*k*_ and **b**_*k*_ are composed of learnable parameters for *k* = 1, …, 4, and *σ* is the ReLU function.

#### Masked locus modeling

Here we describe the loss function of the MLM pre-training task. The objective of the MLM task is to predict the contacts between a set of loci from the input sequence. Given that the [MASK] token will not be used at inference time, it is important that both masked and unmasked loci be included in this set to reduce the mismatch between the pre-training and inference settings. We define 𝒱 ⊆ 𝒯\ ℳ as the set of unmasked loci chosen to enter into the loss. The set of all tokens entering into the loss is then ℐ =ℳ ∪ 𝒱. We found that because 𝒯 already represents such a small portion of the Hi-C matrix, the maximal selection 𝒱 =𝒯\ ℳ helps to accelerate training convergence.

In MLM, we seek to predict the contacts between every pair of loci indexed by ℐ. The index set of the pairwise contacts is 𝒫:= ℐ × ℐ. The loss ℒ is the mean squared error defined as

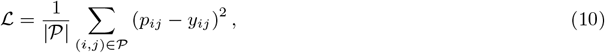

where |𝒫| is the cardinality of 𝒫, the predictions *y*_*ij*_ are defined in Eq. (9), and the targets *p*_*ij*_ are defined as

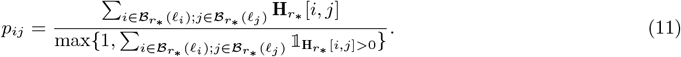

Once again, we take the average over non-zero entries to avoid biasing the loss toward longer loci that have higher total numbers of contacts.

#### Optimization

The pre-training was conducted in two stages. The first stage used a 192-experiment subset of the full dataset and last 6,000 epochs. By reducing the size of the dataset, we were able to train on a single node consisting of 8 NVIDIA H100 GPUs and 1 TB of RAM. In the second stage, we trained over the full dataset using four nodes, each with the same specifications as before, for 5,700 epochs. In both stages, checkpoints were saved at the lowest validation loss, which was evaluated every 50 epochs. Each stage was composed of multiple runs, and each run initialized the model and optimizer from the last saved checkpoint. The hardware usage for pre-training is summarized in Fig. 3.

Optimization was performed using the Adam optimizer^55^ with beta parameters of *β*_1_ = 0.9 and *β*_2_ = 0.98, epsilon parameter *ε* = 10^−9^ and gradients clipped at 1. At the beginning of training, we linearly increased the learning rate *γ* up to 10^−5^ over 500 gradient steps. After this warm-up period, the learning rate was held constant at 10^−5^ for 3,000 steps before decreasing to 10^−6^ over the following 2,000 steps using cosine annealing. Upon reaching 10^−6^, the learning rate was returned to 10^−5^, and the process was repeated in perpetuity with the same period length of 5,000 steps. The optimizer state was saved along with every model checkpoint, so this learning rate schedule was preserved across the two stages. In the final run, we reset the optimizer state and reduced the max learning rate to 10^−6^ following the approach used by large language models^56,57^, but, based on the loss curve, it did not appear to be advantageous in this case. Figures of the learning rate and the training and validation curves are shown in Supplementary Fig. 5.

### Resolution enhancement

As data, we use eight high-coverage Hi-C datasets from Rao et al.^44^, with six in the training set and two in the testing set (Supplementary Table 2). To generate low-coverage input data, we downsample the high-coverage data using a binomial distribution. Let B(*n, p*) represent the binomial distribution, where *n* is the number of trials and *p* is the probability of success. For a downsampling percentage *π*, the low-coverage data 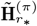 is given as

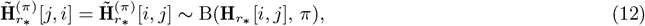

for *i* ≤ *j*. For training, we used *π* ∈ {0.01, 0.1, 1}.

We used a training scheme similar to pre-training, but with four key differences. First, every training sample uses only a single resolution. To complete an epoch, data must be loaded from every experiment at every resolution and at every level of coverage. The set of resolutions remain the same as in pre-training.

Second, the input sequence is formed from two sets of contiguous loci rather than purely randomly. This alteration ensures that a greater percentage of target pixels will lie within the intrachromosomal blocks, which are more data dense and have more complex patterning. The input sequence was formed by first creating two non-overlapping subsequences of length *N* /2. For a given resolution *r*, this was achieved by sampling *i, j* ∼ 𝒰 {0, …, *n*_*r*_ − *N* /2} until | *i* − *j* | ≤ *N* /2. Assuming without loss of generality that *i < j*, we define the index set ℐ = {*i*, …, *i* + *N* /2 − 1, *j*, …, *j* + *N* /2 – 1}. The input loci are then 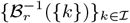.

Third, the target pixels were taken from the high-coverage data regardless of input coverage. Additionally, because every training sample is now at a single resolution, we used target pixels directly from **H**_*r*_ at every resolution, avoiding the approximation errors introduced by pooling after normalization (Eq. (11)). The fourth and final difference is that no masking was used.

Letting 𝒟 be the set of high-coverage experiments, ℛ the set of resolutions, 𝒮 the set of sampling coverages, and 𝒫 = ℐ × ℐ the index set of locus pairs, the resolution enhancement training loss for a full epoch is

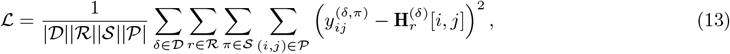

where *y*_*ij*_ are the model predictions and the superscripts denote each variable’s dependence on the data, resolution, and/or coverage. The model was optimized with Adam^55^ for 200 epochs using a batch size of one, and a learning rate of 10^−5^ with a linear warmup for 750 optimization steps. Both the encoder and task head were fine-tuned as this gave better results than only training the task head (Supplementary Figs. 3 and 4).

### Hyperedge identification

Lastly, we considered the problem of hyperedge identification. We formed the Pore-C dataset from GM12878^48^ and BJ and IR fibroblasts^49^. We first created “virtual Hi-C” to use as input to the model. For every hyperedge composed of *K* loci 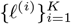, we generated the pairwise edges (*ℓ*^(*i*)^, *ℓ*^(*j*)^) for all *i* ≠ *j, i, j* ≤ *K*. Each edge was tallied and added to an adjacency matrix to create virtual Hi-C.

We considered hyperedges of order 3–5, and preprocessing was performed for each edge size individually. To preprocess, we ranked hyperedges by their observed frequencies and then divided the rank by the total number of unique hyperedges to get normalized frequency values within the unit interval. The normalized frequency of each hyperedge was used as its corresponding weight in the loss function. During training, one negative sample was dynamically generated from each positive sample. To balance the positive and negative samples, the positive weights from each dataset were scaled to have a mean value of one, and all negative weights were set to one. These preprocessing and negative sample generation procedures follow the methodology in^36^.

In this example, we computed the pre-trained embeddings for each dataset at 100 kb lengths. For a *K*-way candidate hyperedge, we selected the embeddings corresponding to the *K* loci and stacked them to form the input sequence **X** ∈ ℝ^*K*×*d*^. The input is pushed through a task head comprised of four transformer encoder layers and a linear output layer. The linear layer maps each embedding to a scalar value, and the scalars are then averaged together to get the predicted logit. Mathematically, the complete task head is given as

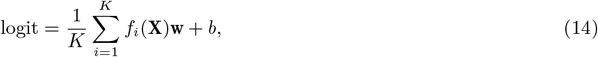

where *f*: ℝ^*K*×*d*^ ↦ ℝ^*d*^ is the *i*th output of the four-layer encoder and the parameters of the linear output layer are **w** ∈ ℝ^*d*^ and *b* ∈ ℝ. The logit can be passed through a sigmoid function to yield a probability *p* = 1/(1 + exp(− logit)).

The training loss was binary cross entropy

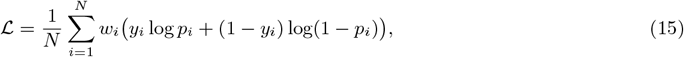

where *y*_*i*_ is one if the loci in the *i*th sample form a hyperedge and zero otherwise, *N* is two times the number of positive samples within a training batch, and *w*_*i*_ are the weights. To optimize, we used Adam^55^ with a batch size of 128 and a learning rate of 10^−5^ for 50 epochs. We used just 100,000 of the positive samples in each dataset for training and reserve the remaining for testing. This amounts to 0.24%, 15.59%, and 38.54% of GM12878, BJ fibroblast, and IR fibroblast datasets, respectively. Once trained, we replaced the virtual Hi-C with real Hi-C to test the model. GM12878 Hi-C comes from Rao et al.^44^, and BJ fibroblast Hi-C from Chen et al.^58^.

We compare to MATCHA^36^ and three multi-correlation methods^45–47^. We set the hidden dimension of MATCHA to be 1,024 to equal that of ARCH3D. The MATCHA architecture uses only node indices as inputs, so it cannot be transferred to a different dataset/cell line without re-training from scratch. Therefore, we trained MATCHA on each dataset separately, using 20% of unique hyperedges from each dataset to train. The three multi-correlation coefficients that we are use are defined as

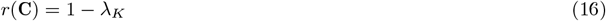

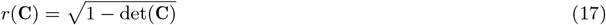

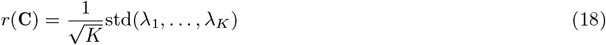

where *λ*_1_ ≥ … ≥ *λ*_*K*_ are the eigenvalues of the correlation matrix **C** ∈ ℝ^*K*×*K*^ and std denotes the standard deviation.

## Data availability

The pre-training corpus is comprised of Hi-C data from the 4DNucleome Program at https://data.4dnucleome.org/ and the ENCODE Project at https://www.encodeproject.org/. The relevant accession numbers can be found in the Supplementary Table 1. For the hyperedge identification experiment, the GM12878 Hi-C is from the 4DNucleome (4DNESLQG7ZKJ) and BJ fibroblast Hi-C from the Gene Expression Omnibus (GEO) (GSE81087). The Pore-C data is available in the GEO for GM12878 (GSE149117) and BJ and IR fibroblasts (GSE211897).

## Code availability

The code is available at https://github.com/ngalioto/ARCH3D.

## Acknowledgments

We thank Ram Prakash, Joshua Pickard, Alvaro Velasquez, and Yannis Kevrekidis for valuable discussions, conversations, and ideas on how to present the results. This work was supported by the Defense Advanced Research Projects Agency (DARPA) award number HR00112490472. Computing resources were provided by an AFOSR DURIP under Program Manager Dr. Fariba Fahroo and grant number FA9550-23-1-006.

## Author information

### Contributions

N.G., C.S., A.G., and I.R. conceived of the study. N.G. designed and implemented the model, conducted the computational experiments, and analyzed the results. N.G. generated the figures in consultation with A.G. and I.R. N.G., A.G., and I.R. wrote and revised the manuscript. C.S. curated the pre-training corpus. A.G. and I.R. provided supervision and secured funding and resources.

## Footnotes

1 https://4dnucleome.org/

2 https://www.encodeproject.org/

