## Supplementary Information for "ARCH3D: A foundation model for global genome architecture"

### Supplementary Notes

#### Supplementary Note 1: Discussion on the input layer

Although our tokenization scheme allows an input locus to be of variable length, the implementation of its numerical representation restricted its format to be a vector in $\mathbb{R}^{n_{r_*}}$ containing contacts with loci of only $r_*$ resolution. In this section, we show how the model can be extended to allow the input loci to be represented as arbitrary-length vectors containing contacts with loci that are of arbitrary resolution. To do this, we first prove the existence of an infinite-dimensional representation of any input locus, allowing us to reformulate the input projection as a functional rather than a matrix. Then, through a discretization method of the user's choice, we can devise a discrete linear projection that is consistent with the underlying infinite-dimensional entities.

To begin, we establish some notation. First, we define the interval $G$ representing the one-dimensional genome. We also define $\mathcal{D}$ to be the $\sigma$-algebra induced by $D$. Together, these form the measurable space $(G, \mathcal{D})$.

Next, we assert that a locus $\mathbf{l}$ can be viewed as the discretization of a measure $\nu_\ell : \mathcal{D} \mapsto \mathbb{R}_{\geq 0}$. Verifying that $\nu_\ell$ meets the definition of a measure is straightforward. In addition to $\nu_\ell$, we can also define the usual Lebesgue measure $\lambda : \mathcal{D} \mapsto \mathbb{R}_{\geq 0}$ over this space. Note that $\nu_\ell$ is absolutely continuous with respect to $\lambda$ for any $\ell \in D$. This can be shown by first noticing that $\lambda(A) > 0$ for any non-empty $A \in \mathcal{D}$. Therefore, $\lambda(A) > 0$ whenever $\nu_\ell(A) > 0$ for every $A \in \mathcal{D}$, satisfying the condition for absolute continuity. Given these properties, the Radon-Nikodym theorem states that there must exist a $\mathcal{D}$-measurable function $f_\ell : G \mapsto \mathbb{R}_{\geq 0}$ such that

$$\nu_\ell(A) = \int_A f_\ell \mathrm{d}\lambda, \quad \forall A \in \mathcal{D}. \tag{1}$$

The implication of this result is that rather than represent each input locus as a vector, the locus can be mathematically represented by the function $f_\ell$. Consequently, we can define a linear projection $\mathcal{A}$ of the input locus as

$$\mathcal{A}(\ell) = \int_G a(x) f_\ell(x) \mathrm{d}x, \tag{2}$$

where $a : G \mapsto \mathbb{R}^d$ maps each point in $G$ into the vector space $\mathbb{R}^d$. Numerically, this integral can be approximated with the Riemann sum

$$\mathcal{A}(\ell) \approx \sum_{i=1}^{N} a(x_i) f_\ell(x_i) \Delta x_i. \tag{3}$$

Notice that this formulation encompasses the fixed discretization $\mathbf{A}\mathbf{l} = \sum_i \mathbf{A}[:, i]\mathbf{l}[i]$ when $\Delta x_i$ corresponds to the Hi-C bin size and the $x_i$ are chosen such that $f_\ell(x_i)\Delta x_i = \ell_i$.

Thus, to evaluate $\mathcal{A}$, we need to find a function $\mathbf{a}$ consistent with the discretized version $\mathbf{A}$ that we have already learned. In other words, we seek a function $\mathbf{a}_\theta$ defined by the learnable parameters $\theta$

$$\theta^* = \arg\min_\theta \sum_j \|\mathbf{a}_\theta(x_j) - \mathbf{A}[:, j]\|_2^2. \tag{4}$$

This formulation can be generalized further by removing the requirement that $\mathcal{A}$ be linear. Theory and methods for learning nonlinear operators can be found in[1–3]. Given that the matrix $\mathbf{A}$, which contains over 588 million learnable parameters, is by far the largest component of ARCH3D, proper choice of parameterization of $\mathcal{A}$ could dramatically reduce the model size, reducing memory requirements and accelerating fine-tuning and inference.

### Supplementary Figures

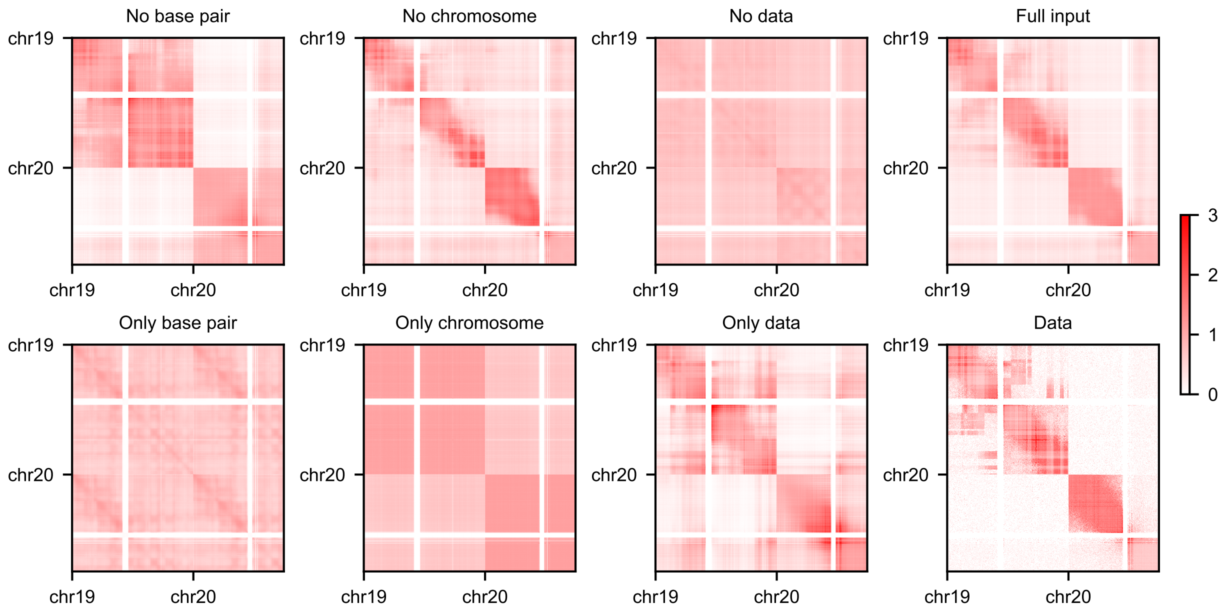


**Supplementary Fig. 1 | Input ablation tests.** The input is composed of a base pair encoding, a chromosome encoding, and the Hi-C data. The reconstruction of chromosome 19 and part of chromosome 20 is shown in 100 kb resolution. The first three plots in the upper row and lower rows show the result of ablating one and two components of the input, respectively. The subplot titled "Full input" shows the reconstruction without ablation, and the one titled "Data" shows the normalized Hi-C data. The base pair and chromosome encodings were ablated by being set to zero. Ablating the data this way, however, would give a trivial result since the model was trained to predict zeros when the data are zero. Instead, the data were ablated by replacing each locus with uniform random noise, scaled such that the sum of every locus is preserved. This figure shows that chromosome encodings help distinguish inter- and intra-chromosomal regions of the Hi-C, base pair encodings allow the model to encode diagonal dominance within intra-chromosomal blocks, and the data enable the model to reconstruct features specific to the Hi-C experiment.

**a**

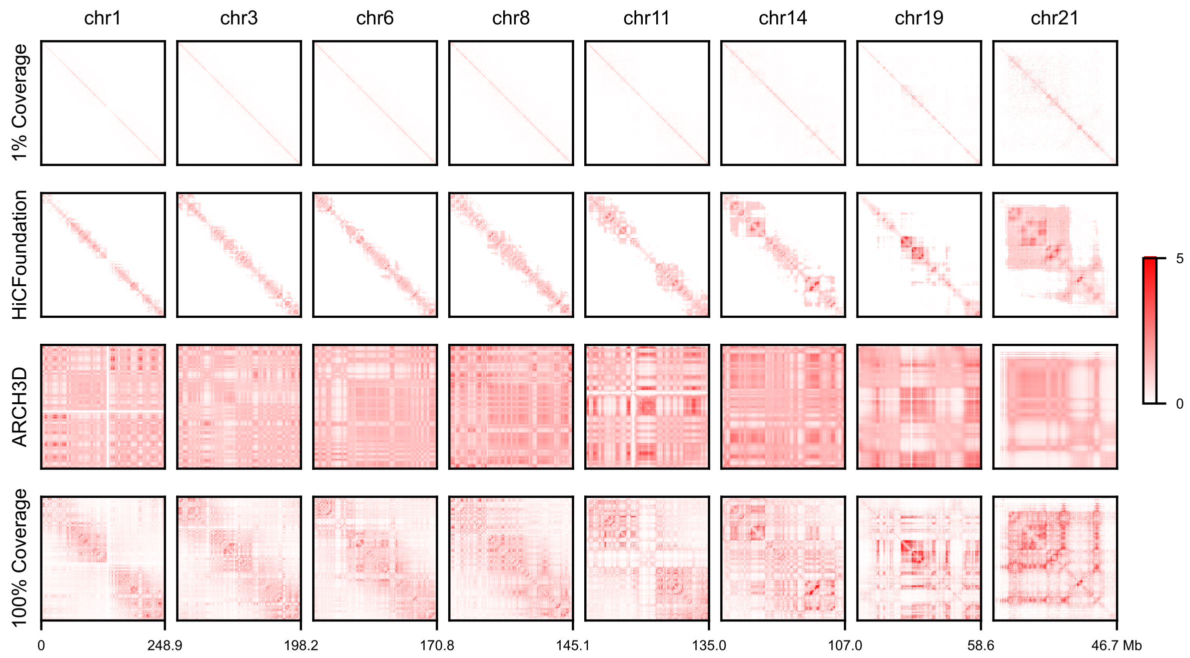


**b**

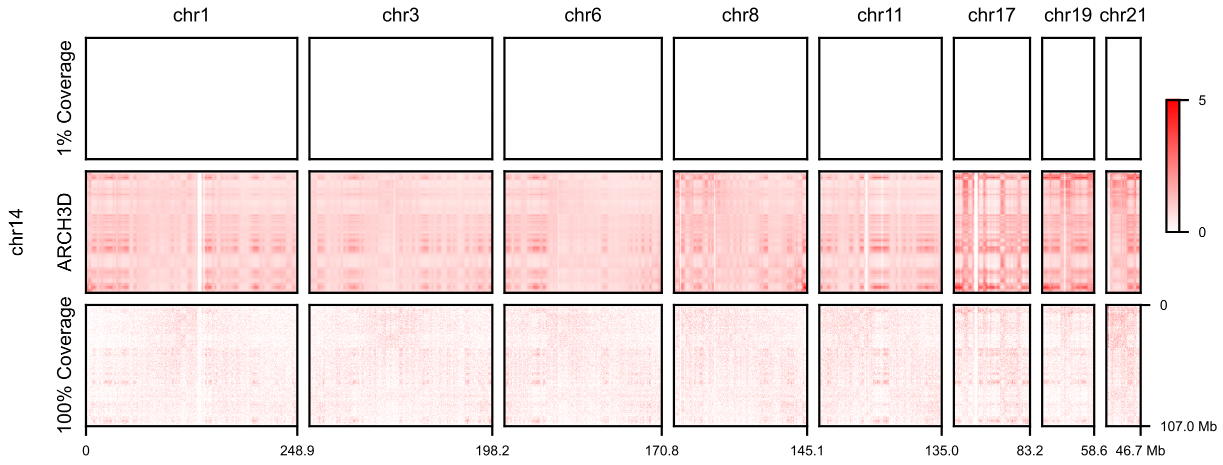


**Supplementary Fig. 2** | Resolution enhancement on HUVEC at 100 kb resolution from 1% coverage (≈1M counts). **(a)** Prediction of intrachromosomal contacts. At this sparsity level, HiCFoundation[4] begins to degrade and cannot even fill its 22.4 Mb sliding window along the diagonal. ARCH3D predictions fill the full contact map and still capture many of the high-level compartments. **(b)** Prediction of interchromosomal contacts. Despite the fact that 99.94% of interchromosomal pixels are zero, ARCH3D produces dense contact maps that visually match the features of the full-coverage data.

**a**

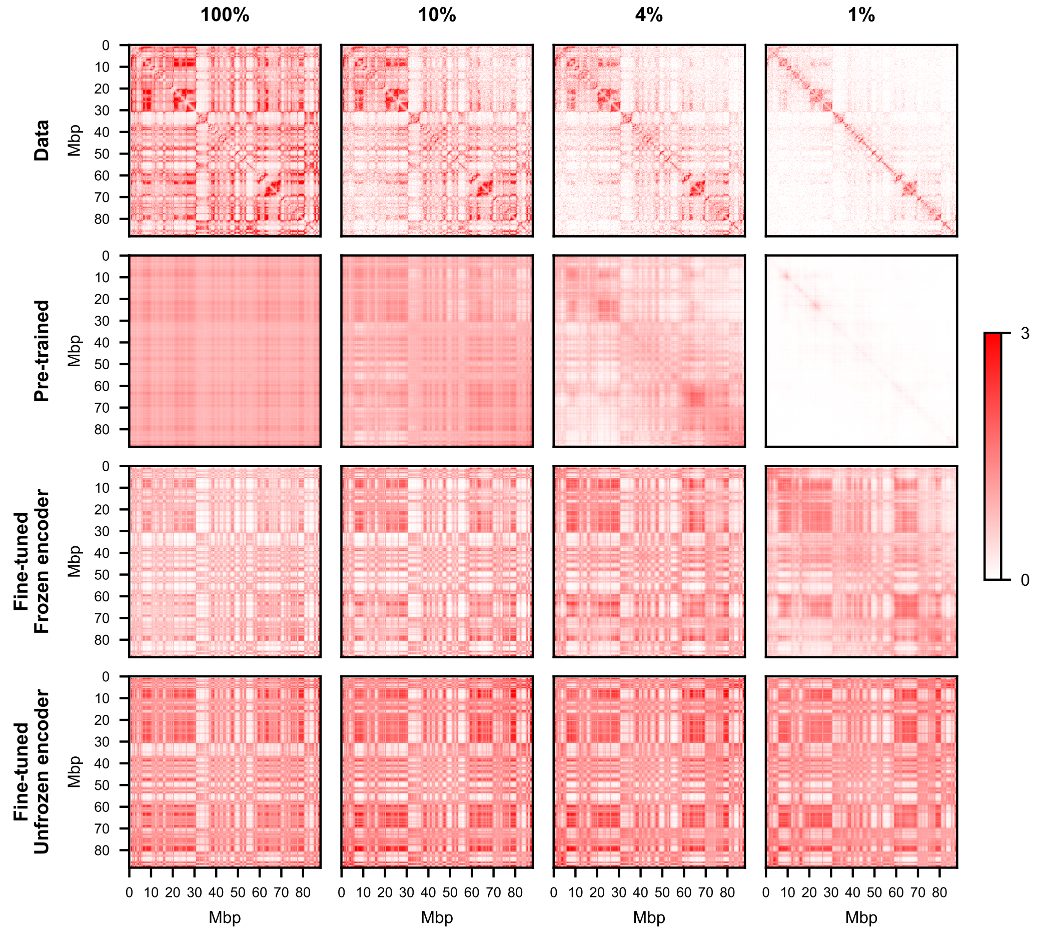


**b**

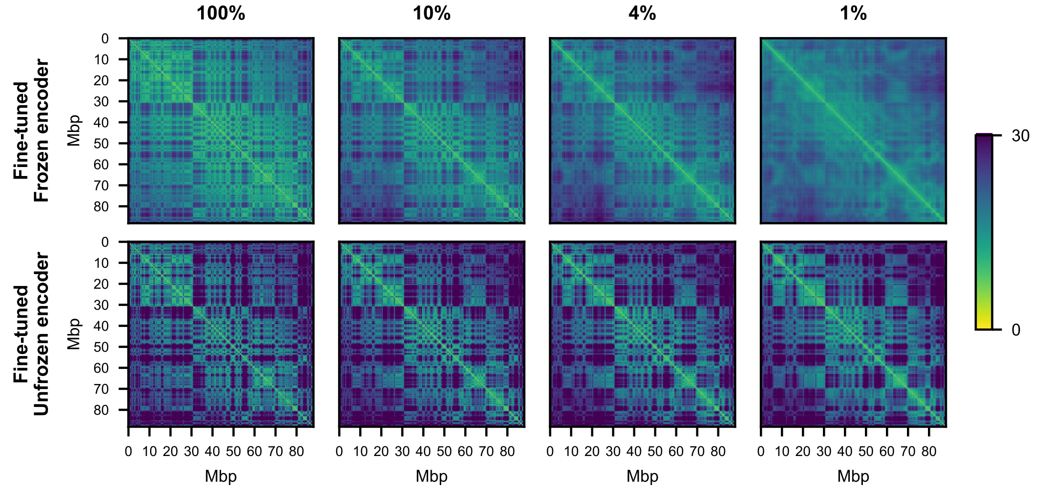


**Supplementary Fig. 3 | Resolution enhancement of IMR-90 under different fine-tuning strategies.** **(a)** Reconstructed pixels. Fine-tuning the task head improves pixel reconstruction, but the best performance comes from fine-tuning both the task head and encoder. **(b)** Euclidean distance between locus embeddings. The upper and lower rows show the embedding distances from the pre-trained and fine-tuned encoders, respectively. After fine-tuning, the embeddings are not as sensitive to the Hi-C coverage.

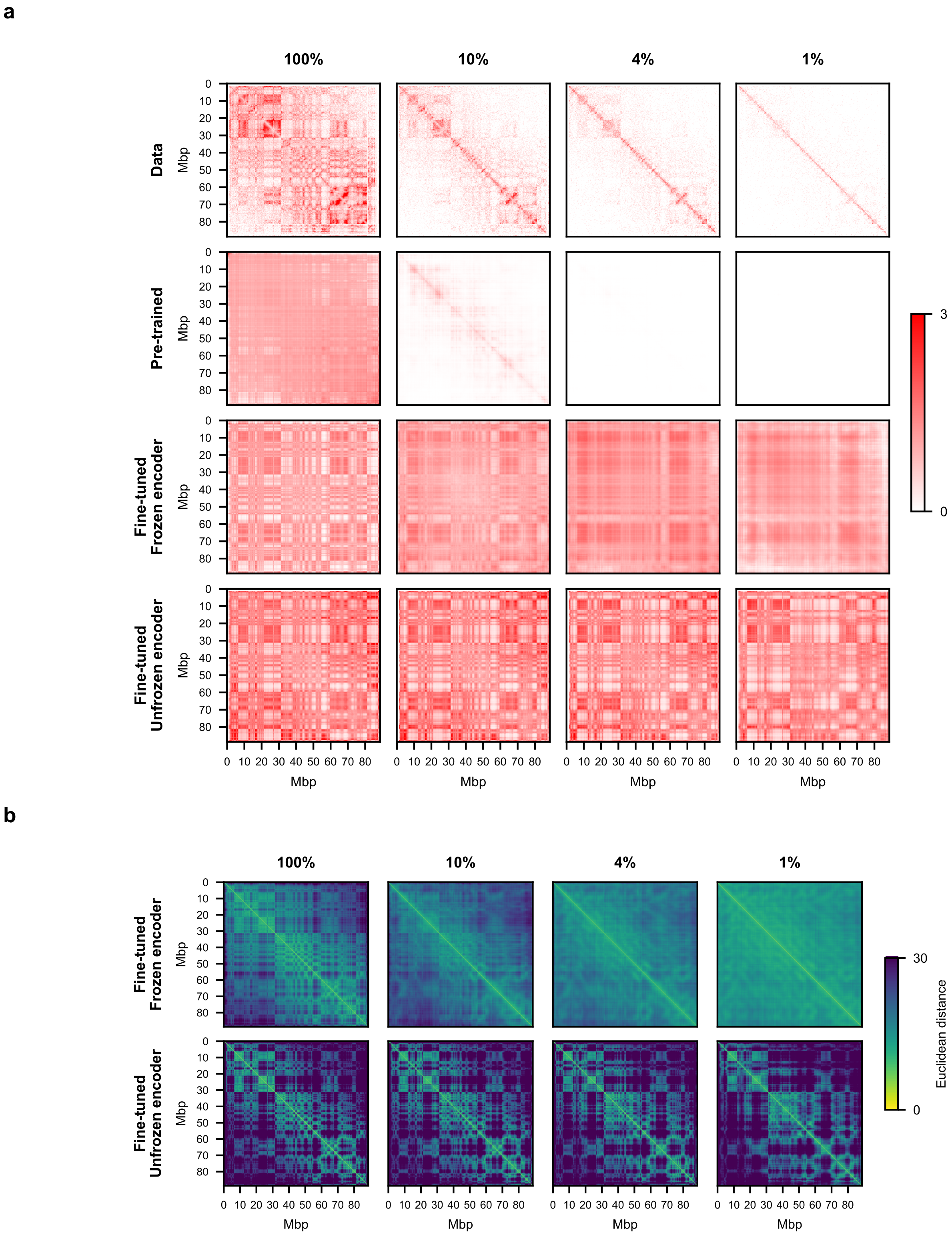


**Supplementary Fig. 4 | Resolution enhancement of HUVEC under different fine-tuning strategies. (a)** Reconstructed pixels. Without fine-tuning, the model simply predicts zeros at 4% and 1% coverage. Fine-tuning results in significant increase in coverage of ARCH3D predictions. **(b)** Euclidean distance between locus embeddings. Similarly to Supplementary Fig. 3, fine-tuning results in embeddings that are less sensitive to the Hi-C coverage.

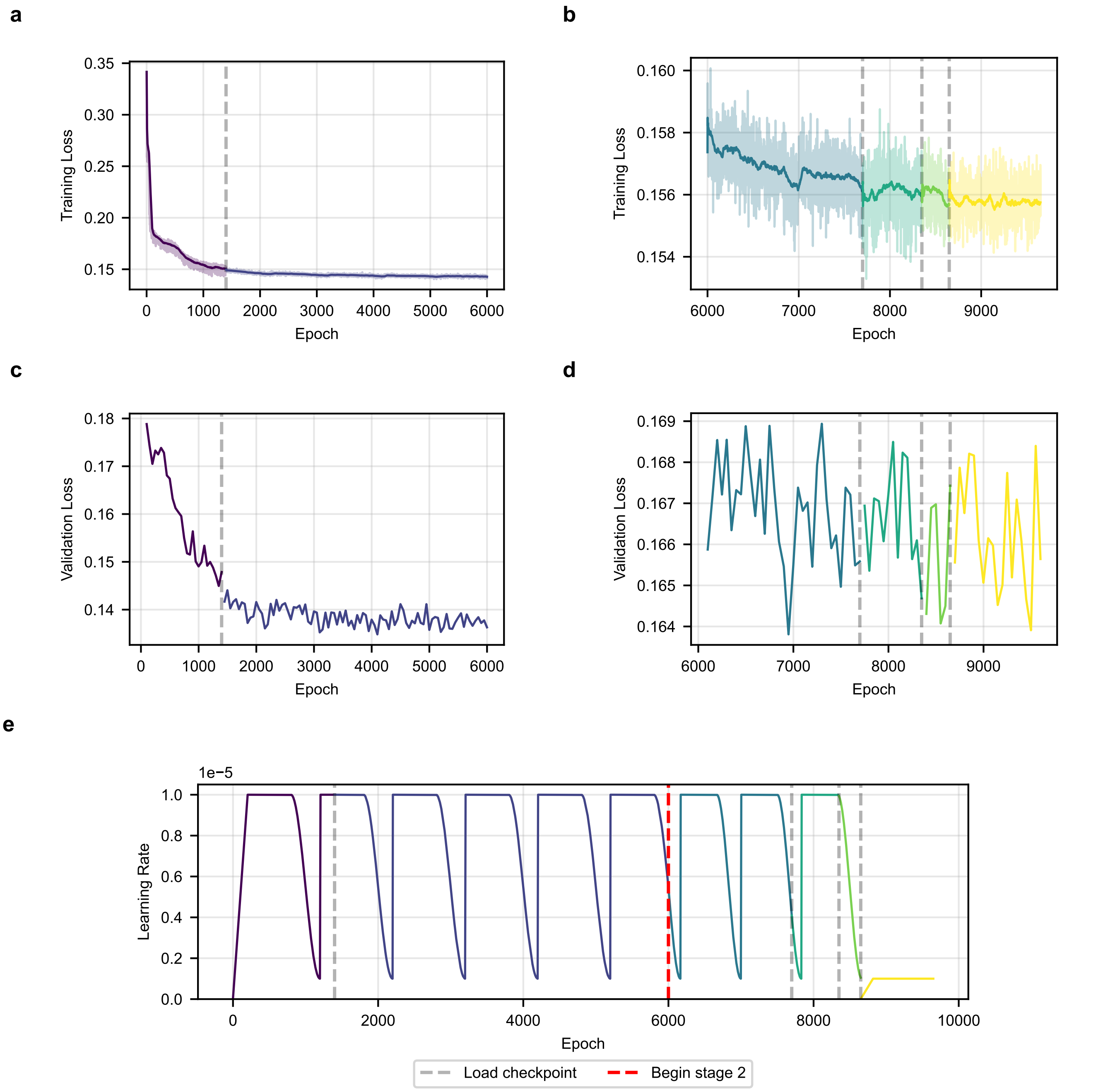


**Supplementary Fig. 5 | Pre-training metrics** (see Table 3). **(a), (b)** Training loss in stages 1 and 2. The dark line shows the moving average over a window of 50 epochs, and the faded lines show the per-epoch loss values. **(c), (d)** Validation loss in stages 1 and 2. The validation loss is computed every 50 epochs. **(e)** Learning rate over time. The learning rate begins with a linear warmup over the first 1,000 optimization steps to a maximum value of $10^{-5}$. The learning rate is held constant for 3,000 steps, then reduced to $10^{-6}$ via cosine annealing over 2,000 steps. The learning rate is then returned to the maximum value and the cycle repeats. In the final training segment, the optimizer is restarted, and the maximum learning rate is reduced to 10% of its initial value, following other approaches[5,6]. Although the learning rate progresses according to optimization steps, we chose to plot it here against epochs to align with the loss plots (**a**)–(**d**). As a result, the cycling period appears shorter in stage 2.
